# Single-cell transcriptomic profiling of *bam* mutant tumor reveals germline heterogeneity and *gcrf1* as a modulator in *Drosophila* germ cells

**DOI:** 10.1101/2025.10.14.682265

**Authors:** Zhipeng Sun, Yujun Zeng, Todd G Nystul, Guohua Zhong

**Author notes:** These authors contribute equally to this work (Zhipeng Sun, Yujun Zeng). Corresponding author (G. Zhong). (N.T.).

## Abstract

The *bam* mutant ovary of *Drosophila* represents a classic tumor model caused by germline stem cell (GSC) differentiation defects. To date, its molecular and genetic features have rarely been characterized in detail at the single-cell resolution. Here, we performed single-cell RNA sequencing (scRNA-seq) to comprehensively delineate the transcriptomic landscape and identify distinct germline cell types in *bam* mutant ovaries by using *in situ* hybridization. Differentially expressed gene analysis and PAGA plots reveal different transcriptional profiles and developmental relationships in ovarian cells. Based on the expression pattern of *eggpl*, a useful marker for undifferentiated germ cell identity, and morphological differences in *bam* mutant ovarioles, two potentially distinct germ cell states are distinguished. Comparative single-cell analysis reveals the potential regulatory network and cellular communication in subclusters of undifferentiated germ cells, and contributes to the identification of *gcrf1* as a novel marker gene for female GSC, which involves in the regulation of early germ cell proliferation and *Drosophila* fertility. Collectively, our study not only provides insights into tumorigenesis caused by GSC differentiation defects but also offers a valuable transcriptomic resource that can be mined for the reproductive features of *bam* mutant tumors by community.

## Introduction

Tissue homeostasis and development depend on tight regulation of stem cell proliferation and differentiation. Under normal conditions, adult stem cells undergo continuous self-renewal and differentiation in a controlled manner to replenish damaged cells and dead cells caused by aging, injury or disease (1–3). Ovarian germline stem cells (GSCs) in *Drosophila melanogaster* experience constant division throughout adult lifespan to sustain oocyte production, making them an excellent *in vivo* model for stem cell research (4, 5). Located at the anterior tip of ovariole within the germarium, each GSC divides asymmetrically: one daughter cell remains anchored to the niche to preserve stem cell fate, while the other daughter cell, known as the cystoblast (CB), is displaced from niche and commits to differentiation (6). The CB undergoes four rounds of mitosis with incomplete cytokinesis, generating a 16-cell interconnected germline cyst linked by ring canals and an actin-based branched fusome that passes through ring canals from one cell to the next (7, 8).

As a key ubiquitin associated protein, *bag-of-marbles* (*bam*) plays a crucial role in promoting the specialized transition of GSC to CB and early cysts (9). In GSCs, *bam* expression is transcriptionally silenced by a complex of pMad and Med to maintain stem cell identity (10, 11). Upon GSC division, *bam* is upregulated in the CB where it promotes differentiation by forming a complex with the RNA-binding protein benign gonial cell neoplasm (Bgcn) and Sex lethal (Sxl) to repress the translation of self-renewal factors such as *nanos, pumilio* and *eIF4a* (12). Additionally, Bam competes with components of the COP9 signalosome, particularly Csn4, to modulate the activity of the COP9 complex, thereby influencing GSC self-renewal and differentiation (13, 14). Disruptions of this balance between GSC proliferation and differentiation by genetic mutations or external cues can lead to severe germline defects or sterility. For example, ectopic expression of *bam* in early germ cells leads to complete GSC loss and an empty ovariole phenotype (10), while loss of *bam* function blocks CB differentiation and results in the formation of undifferentiated germ cell tumors (15, 16). However, few studies have demonstrated the molecular signatures or cellular composition of *bam* mutant tumors.

Single-cell RNA sequencing (scRNA-seq) has emerged as a powerful tool to resolve cellular heterogeneity in different tumor tissues such as basal cell carcinoma, small-cell neuroendocrine cervical carcinoma and non-small cell lung cancer, enabling the detailed dissection of tumor composition, cell-cell interaction, and the tumor microenvironment at unprecedented resolution (17–19). In *Drosophila* tumor models, scRNA-seq has been instrumental in elucidating how genetic mutations affecting apicobasal polarity or tumor suppressors like *Rbf* drive epithelial multilayering and invasive behavior (20). By using scRNA-seq, researchers have verified that loss of polarity genes (e.g., *l*(*2*)*gl*) induces cellular plasticity, which is mediated by Keap1-Nrf2 signaling in promoting multilayer formation, and activates lineage-specific transcriptional programs (21). In the follicular epithelium, polarity loss combined with oncogenic cues, like Notch overexpression, reveals a context-dependent responsiveness to JAK/STAT signaling pathway, contributing to phenotypic heterogeneity (22). These findings provide a single-cell view of tumor cell states and identify previously unrecognized pathways linking polarity loss to early tumor progression.

In our study, we employed scRNA-seq to systematically characterize the cellular composition and transcriptional landscape of *bam* mutant ovaries. Through differential gene expression analysis and pseudotime inference, we uncovered distinct transcriptional states and developmental trajectories in the germ cell lineage. Notably, we identified two germ cell states that differ in the levels of *eggpl* expression and tumor morphology, revealing previously unrecognized heterogeneity within the undifferentiated germ cell compartment. Furthermore, comparative analysis revealed potential regulatory networks and cellular communication pathways among subclusters, and discovered a novel GSC marker, *CG13949*, that modulates early germ cell proliferation and fertility, which we have named *germ cell renewal factor 1* (*gcrf1*). Together, these findings provide new insights into the cellular and molecular basis of GSC-derived tumorigenesis and establish a valuable transcriptomic resource for future studies of *bam* mutant ovarian biology.

## Materials and methods

### Fly stocks and maintenance

The following stocks were used in this study: *y^1^w^1118^*, *nos*-Gal4, *bam^Δ86^/*TM3 (BDSC 5427), *eggpl*::GFP;*nos*-Gal4, UAS-*bam*-RNAi (BDSC 58178), UAS-*gcrf1*-RNAi (BDSC 61207), *AstC-R2* T2A-Gal4 and UAS-mCD8::GFP were gifts from Ryusuke Niwa’s lab (Life Science Center for Survival Dynamics, Tsukuba Advanced Research Alliance (TARA), University of Tsukuba, Japan). Notably, *gcrf1* knock-in and UAS-*gcrf1*-GFP lines were constructed according to standard protocols which applied in our previous work (23).

All fly stocks and crosses were maintained at 24 ℃ in artificial climate chamber and fed on fresh standard cornmeal-agar medium.

### Single-cell suspension preparation, scRNA sequencing and clustering

300 fresh ovary tissues from adult homozygous *bam^Δ86^* flies were dissected under stereomicroscope (Leica, SAPO, Germany). After collection, the tissues were rinsed by PBS buffer 3 times and dissociated by adding 70 μl of 5% (w/v) trypsin (Invitrogen, cat. no. 27250-018) and 70 μl of 2.5% (w/v) collagenase (Invitrogen, cat. no. 17018029) to 560 μl of PBS, and incubated with a shaking speed of 60 r.p.m for 15 min. The cell suspension was filtered using a 40-μm nylon cell strainer (Falcon, Cat. no. 352340) and centrifuged for 5 min at 425 × *g*, 4°C. The pellets were resuspended with 200 μl of serum-free Schneider’s insect medium, and the cell viability was examined by using 0.4% trypan blue (Solarbio, cat. no. T8070). The concentration of cell suspension was diluted to about 1×10^6^ cells/ml for scRNA-seq.

Single-cell libraries were constructed using chromium single-cell 3’ Library (v2) kit via End Repair, A-tailing, Adaptor Ligation, and PCR according to the manufacturer’s protocol and sequenced on Illumina HiSeq 4000 with a custom paired-end sequencing mode 26 bp (read 1) × 98 bp (read 2). All reads were processed using Cell Ranger Single Cell Software Suite (v6.1; http://software.10xgenomics.com/single-cell/overview/welcome) with the default parameters. FASTQs files were aligned to the *Drosophila* reference genome (dm6; https://www.ncbi.nlm.nih.gov/assembly/GCF_000001215.4#/st) by STAR RNA-Seq aligner. Next, a gene-barcode matrix was generated by counting unique molecular identifiers (UMI) and filtering non-cell associated barcode, and it was imported into the Seurat (v4.0.4) R toolkit for quality control following the criteria: (1) gene counts > 3000 per cell; (2) UMI counts > 12,000 per cell; and (3) percentage of mitochondrial genes > 30%. We finally obtained 11,561 out of 15,520 cells with 8,022 median UMIs per cell and 2,307 median genes per cell for downstream analysis.

For the clustering, the principal component analysis (PCA) was applied to normalize and filter the gene-barcode matrix and to reduce feature dimensions. The top five major components were selected to obtain the visualized 2D clustering image using T-distributed stochastic neighbor embedding (tSNE). The graph-based clustering method was used to categorize cells with similar expression patterns of marker genes into the same cluster. The 11 unsupervised categories using the default resolution parameter (*R* = 0.5) were presented.

### Partition-based graph abstraction (PAGA) analysis

The developmental trajectory analysis was performed by using PAGA in this study. Based on our single-cell RNA dataset, an PAGA graph was generated by quantifying the connectivity of distinct clusters. The computations and algorithms were performed at https://scanpy.readthedocs.io/en/stable/index.html.

### Gene Ontology (GO) term and KEGG pathway enrichment analysis

Gene Ontology (GO) term analysis was performed to understand the highly expressed genes that correspond to biological process, molecular function and cellular component. The peak-related genes were mapped to GO terms in the GO database (http://www.geneontology.org/), and the significantly enriched GO terms were defined by a hypergeometric test. KEGG enrichment analysis identified significantly enriched metabolic pathway and signal transduction pathways in differentially expressed genes.

### Generation of Sankey plot

The Sankey plot was and visualized through the. To visualize the correspondence of cell populations between two scRNA-seq datasets, we first preprocessed and normalized the datasets using standard workflows in Seurat (v4.0.4) R. The cluster annotations were obtained from prior unsupervised clustering and marker-based classification. A contingency matrix was then generated to quantify the overlap in cell-type identities across the two datasets. This matrix was used to compute the frequency of transitions between clusters. The resulting transition frequencies were generated using the networkD3 package and visualized using the sankeyNetwork function, illustrating the proportional relationship of cell populations.

### SCENIC analysis

Gene regulatory network analysis was conducted by using SCENIC v1.3.1 and Rcis Target v1.16.1. The Rcis Target package was used to analyze excess transcription factor (TF) binding motifs, and the Spearman correlation between TFs and target genes was calculated by the GENIE3 function.

### Cellular communication analysis

According to the gene expression matrix and clustering information, the CellPhoneDB V5 software was used to predict the abundant ligand-receptor interaction among distinct clusters, and analyzed the number of ligand-receptor pairs and expression abundance in each cluster (24). CellChat V2.1.0 software was used to analyze the communication ligand-receptor pairs between cell subpopulations and construct interaction network at ligand-receptor level (25)

### Construction of phylogenetic tree on Gcrf1 protein and protein-protein interaction (PPI) network prediction

To identify homologous protein with high amino acid sequence similarity (>50%) to Gcrf1, online BLAST analyses were performed using the NCBI database (https://www.ncbi.nlm.nih.gov/). The resulting sequences were then used for phylogenetic tree construction and visualization via the Interactive Tree of Life (iTOL) online tool (https://itol.embl.de/).

The PPI network was produced by entering the amino acid sequence of Gcrf1 and modifying the outcome in online platform (https://fgrtools.hms.harvard.edu/MIST/).

### RNA *in situ* hybridization

The digoxigenin (DIG)-labeled probe was synthesized using wild-type *Drosophila* DNA as a template, and amplified the exon regions of *gcrf1* by using the primers with SP6 sequence (ATTTAGGTGACACTATAGAAGNG) according to the instruction of KAPA HiFi PCR Kit (Roche Diagnostics, cat. no. 07958927001). The primers were listed in **Supplementary Table S1**.

The *in situ* hybridization was performed according to standard protocol described in our previous work (23).

### Immunofluorescence staining and confocal imaging

Adult ovaries were dissected in PBS and fixed in 4% paraformaldehyde fix solution (PFA), 0.1 M Hepes, PH 7.4 for 30 min with rotation. Then the tissues were washed 3 × 15 min with 500 μl 0.1% PBT (0.1% Triton X-100 in PBS), blocked in 5% NGS (5% normal goat serum in 0.1% PBT) for 1 h and incubated with primary antibody overnight at room temperature. The following day, the tissues were washed 3 × 15 min with 500 μl PBT and incubated with secondary antibody for 3 h with rotation. The Hoechst 33258 (Sigma-Aldrich, cat. no. 23491454) was used to label the nucleus at final step. Finally, the tissues were mounted on slides in Vectashield mounting medium and stored at 4°C. For pMad staining, the samples were recommended to fix in 4% PFA for 50 min and washed the tissues with 0.1% PBT 3 times for 3 h at least.

Primary and secondary antibodies used in our study were: rabbit anti-α-Spectrin (1:100; Developmental Studies Hybridoma Band (DSHB)), rabbit anti-pMad (1:800; Cell Signaling), mouse anti-HA (1:1,000; Abmart). Alexa Fluor 488 and 555 conjugated goat secondary antibodies (gifts from Yu. Cai, Temasek Life Sciences Laboratory, Singapore) against mouse (1:500) and rabbit (1:1,000).

### Bioassay

Ovary morphological analysis. The ovary phenotype of control, *gcrf1* RNAi and *gcrf1* overexpression lines was observed, and the images were captured by stereomicroscope (Leica, S APO). The ovary size was measured by using Image J V1.8.0 software.

Oviposition. 10 pairs of adult flies were fed in fly vials (cat: 51-0800, Biologix) supplementing with fresh food for 10 days. The number of eggs laid on the medium surface was calculated on a daily basis.

### Quantitative real-time PCR (qRT-PCR) analysis

Total RNA was extracted from 100 mg ovary with RNAiso Plus (TaKaRa, Japan), and the cDNA was synthesized using PrimeScript™ RT reagent Kit with gDNA Eraser (Perfect Real Time) according to the instructions (Takara Biomedical Technology, Beijing, China). Then, real-time PCR was carried out with iTaq™ SYBR® Green Super mix (Bio-Rad, Hercules, CA, USA), gene primers were designed with Primer 5.0 software (**Supplementary Table S2**), and qPCR reactions were conducted on CFX96 System (Bio-Rad) with following conditions: initial denaturation at 95℃ for 30 s, 40 cycles of denaturation at 95℃ for 15 s, annealing at 55-60℃ for 20 s, elongation at 72℃ for 20 s.

### Statistical analysis

The statistical analyses were conducted by Wilcoxon rank sum test which performed via the SAS statistical software package version 8.1 (Microsoft, USA). All graphical images were produced by using GraphPad Prism software package version

9.5 for windows (Microsoft, USA).

## Results

### The global cellular landscape of *bam* mutant ovarioles

Immunofluorescence imaging of *bam^Δ86^* mutant ovarioles reveals a large number of GSCs and GSC-like cells, which can be distinguished by labeling with anti-pMad and anti-α-spectrin antibodies, as well as a variety of somatic cells, including TF cells, cap cells, escort cells, follicle stem cells and differentiated follicle cells (Fig. 1A). To investigate the transcriptional profiles of *bam^Δ86^* ovarian tumors at single-cell resolution, we performed scRNA-seq analyses. We obtained a total of 11,561 highquality cells after standard quality control and filtering steps, and we used the standard Seurat pipeline to categorize the cells into 11 distinct clusters (Fig. 1B). We were able to assign preliminary annotations to ten of these clusters based on differential expression of canonical markers. Five clusters expressed the germ cell specific marker *vas*, and were thus identified as germ cell clusters. We designated one of these clusters as GSC & GSC-like, based on its strong expression of *vas* and *HP6* (26), which is a marker for GSCs and early cyst cells, and another of these clusters as Undifferentiated Germ Cell-1, based on its moderate levels of *vas* and *chinmo* expression (23, 36). In addition, we designated a third cluster as Undifferentiated Germ Cell-2 based on its strong expression of *HP6*, *bam* and *mnd* (26), the fourth cluster, which strongly expresses *chinmo*, as Differentiated Germ Cell, and the last cluster, which strongly expresses *CG15628* (26, 27), as Older Germ Cell. Likewise, we annotated the somatic clusters based on differential expression of canonical markers. Specifically, we identified a TF & Cap Cell cluster that expresses *tj* and *Lmx1a* (26), an Escort Cell & FSC cluster that expresses *tj* and *GstS1* (23, 36), a Terminal Follicle Cell cluster that expresses *ct*, a Main Body Follicle Cell cluster that expresses *Vm26Aa* (27), and a Muscle Cell cluster that expresses *Neurochondrin* (30) (Fig. 1C). In addition, we subclustered the TF cells and Cap Cells into separate clusters. Although no specific marker genes were discovered in the Unknown cluster, we found that the upregulated differentially expressed genes were primarily enriched in DNA replication, RNA transport, nucleotide excision repair, oxidative phosphorylation, pyrimidine metabolism and metabolic pathways, implicating a mixture of mitotic follicle cells and endocycling follicle cells in Unknown cluster (Fig. 1D and Fig. S1 B).

**Fig. 1.**
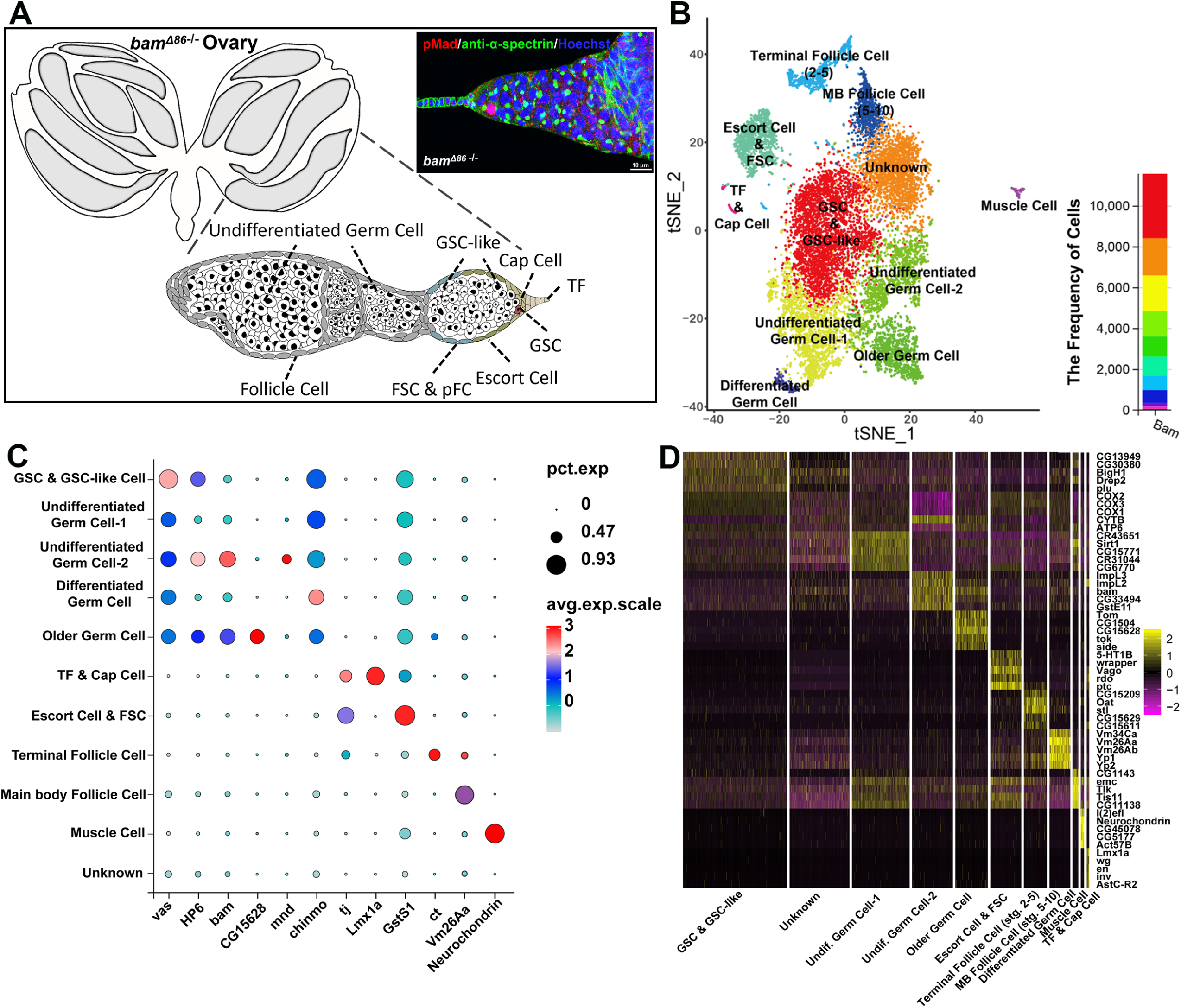
Construction of single-cell transcriptomic atlas in *bam* mutant ovary. **(A)** Illustration of general ovary, typical ovariole and germ cell lineages marked by pMad (red), α-spectrin (green) and Hoechst (blue) in *Drosophila bam^Δ86−/−^*ovary. **(B)** t-distributed scholastic neighbor embedding (t-SNE) plot of 11 distinct clusters determined by cell types and their proportion. **(C)** Dot plot shows the gene expression patterns of canonical markers for germ cells and somatic cells. Dot size indicates the number of cells expressing the marker genes. **(D)** Heatmap of top enriched 5 cell-type markers genes in 11 major cell types. Each column represents a cell, and purple color indicates low gene expression level, while the yellow color indicates high gene expression level.

### Distribution of marker expression and features in somatic cells

Two representative markers were selected from the differentially expressed genes to demonstrate the distribution of marker expression in four somatic cell clusters (Fig. 2A). DEG analysis of the TF and Cap cells identified a G-protein coupled receptor, *AstC-R2*, as a novel marker for a subset of TF cells and we validated this with a GFP-tagged allele of *AstC-R2* (Fig. S2A-B). KEGG enrichment of upregulated genes revealed that several key developmental pathways such as MAPK, Hippo, AGE-RAGE signaling pathways are upregulated in TF& Cap Cell. The pathways enriched in Escort Cell & FSC and Terminal Follicle Cell (stg.2-5) are related to the phagosome, biosynthesis of amino acids, carbon metabolism, rheumatoid arthritis and focal adhesion, while only genes related to the ribosome were significantly upregulated in MB Follicle Cell (stg. 5-10) (Fig. S1A and Fig. 2B). The cell type differentiation trajectory of somatic cells was reflected by PAGA plot abstraction showing that TF & Cap Cell cluster flowed to Escort Cell & FSC and Terminal Follicle Cell (stg.2-5) flowed to MB Follicle Cell (stg. 5-10) (Fig. 2C-D). In addition, the PAGA graph predicted a strong connection between TF & Cap Cell and Escort Cell & FSC, and high relation between Terminal Follicle Cell (stg.2-5) and MB Follicle Cell (stg. 5-10) (Fig. 2E). We found fewer follicle cell types in the *bam* mutant dataset than have been reported previously in wild-type ovary atlases (26, 27, 36), which is consistent with the observation that egg chamber development is hindered by the abnormal germ cell differentiation in *bam* mutant ovarioles (15, 16).

**Fig. 2.**
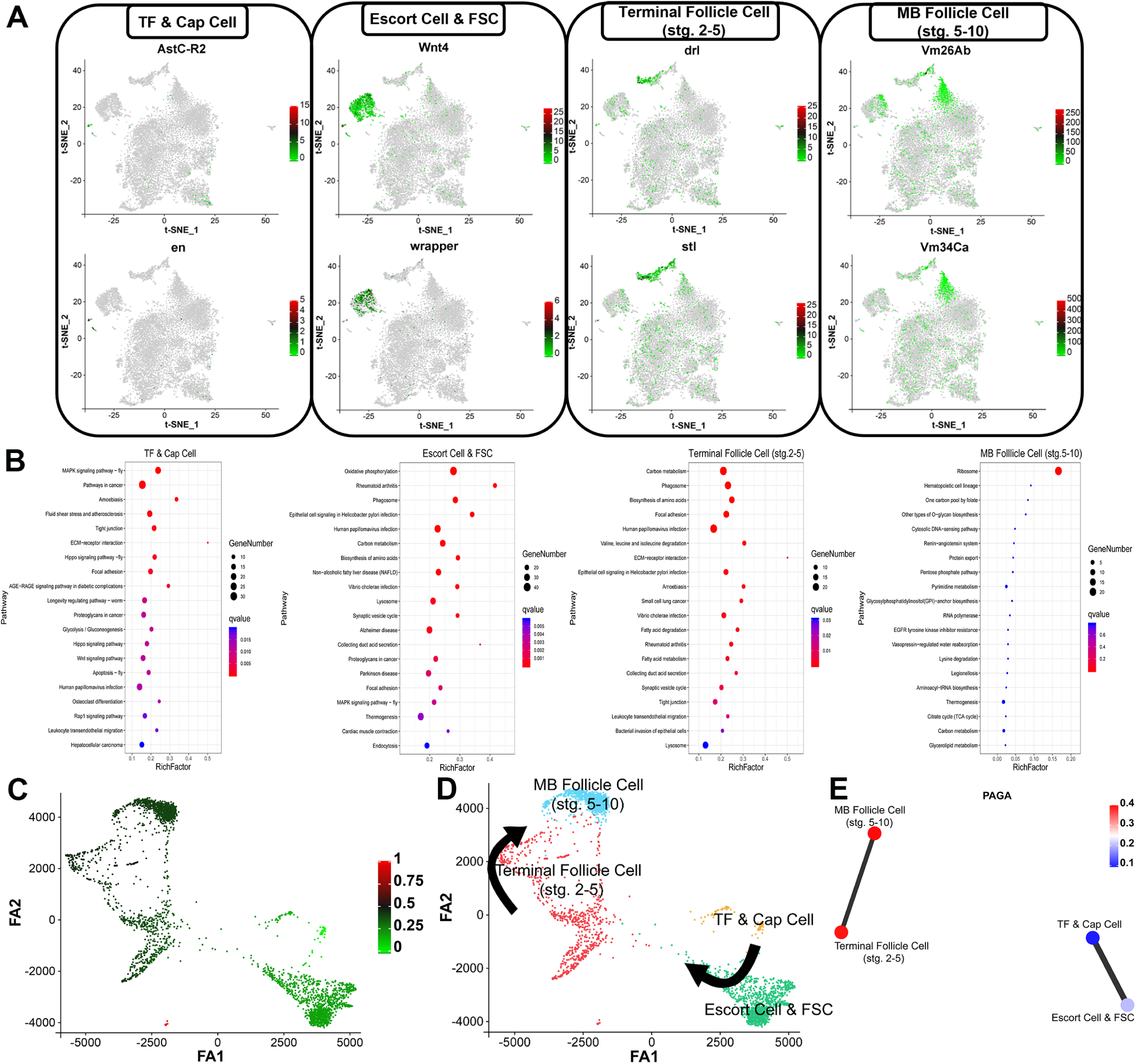
Identification and cellular connectivity of somatic cell types. **(A)** Feature plots on tSNE shows the canonical marker expression used to annotate distinct cell types of somatic cell clusters in *bam* mutant ovary. **(B)** KEGG analysis on top 20 pathways of upregulated genes in 4 somatic cell clusters **(C)** PAGA-initialized single-cell embedding shows the predicted developmental trajectories of somatic cell lineage. The light green dots indicate the cells at early stage, while the dark green dots indicate the cells at late stage. **(D)** Annotation of cell types in PAGA-initialized single-cell embedding trajectories. **(E)** PAGA graph for visualization of cell type interaction between TF & Cap Cell/Terminal Follicle Cell (stg. 2-5) and Escort Cell & FSC/MB Follicle Cell (stg. 5-10).

### Identification of transcriptional profiles and two types of developmental states in *bam* mutant germline

To further characterize gene expression profiles specific to distinct germ cell clusters, we selected candidate markers that are exclusive or highly expressed in target cell clusters on tSNE plot and visualized several maker expression patterns by using *in situ* hybridization. We found enrichment for expression of *gcrf1* in GSC & GSC-like cluster, *CG43293* in Undifferentiated Germ Cell-1 cluster, *CG10802* and *mnd* in Undifferentiated Germ Cell-2 cluster, *CG1143* and *CG34232* in Differentiated Germ Cell cluster, and *CG15628* in Older Germ Cell cluster (Fig. 3A). Comparison of germ cell composition between wild-type and *bam*-mutant ovaries revealed that GSCs primarily differentiate into nurse cells and oocytes in wild type, while in the *bam* mutant, GSCs and their progeny are largely redirected into expended pools of undifferentiated germ cells, differentiated germ cell and older germ cells. Consistent with the original description of *bam* mutant tumors (28), this result confirms that loss of *bam* leads to an accumulation of GSC-like cells and impaired transition toward normal differentiated states (Fig. S3A). The enriched KEGG-pathway-related terms indicated an expression bias for DNA replication, cell cycle, mismatch repair and RNA transport associated genes in GSC & GSC-like cluster, which is quite similar to the previous findings in wild-type ovary scRNA-seq studies (23, 27). In other *bam* mutant germline clusters, the up-regulated expression genes involving in hepatitis B, erbB signaling pathway, viral carcinogenesis and hippo signaling pathway were enriched in both Undifferentiated Germ Cell-1 cluster and Differentiated Germ Cell cluster, whereas the Undifferentiated Germ Cell-2 cluster has an enrichment of genes in aminoacyl-tRNA biosynthesis, protein processing in endoplasmic reticulum and RNA transport and degradation. An enrichment of foxO signaling pathway apoptosis and TNF signaling pathway in Older Germ Cell cluster suggested a potential activation of stress response and programmed cell death mechanisms during late-stage germ cell development (Fig. S1A and Fig 3B).

**Fig. 3.**
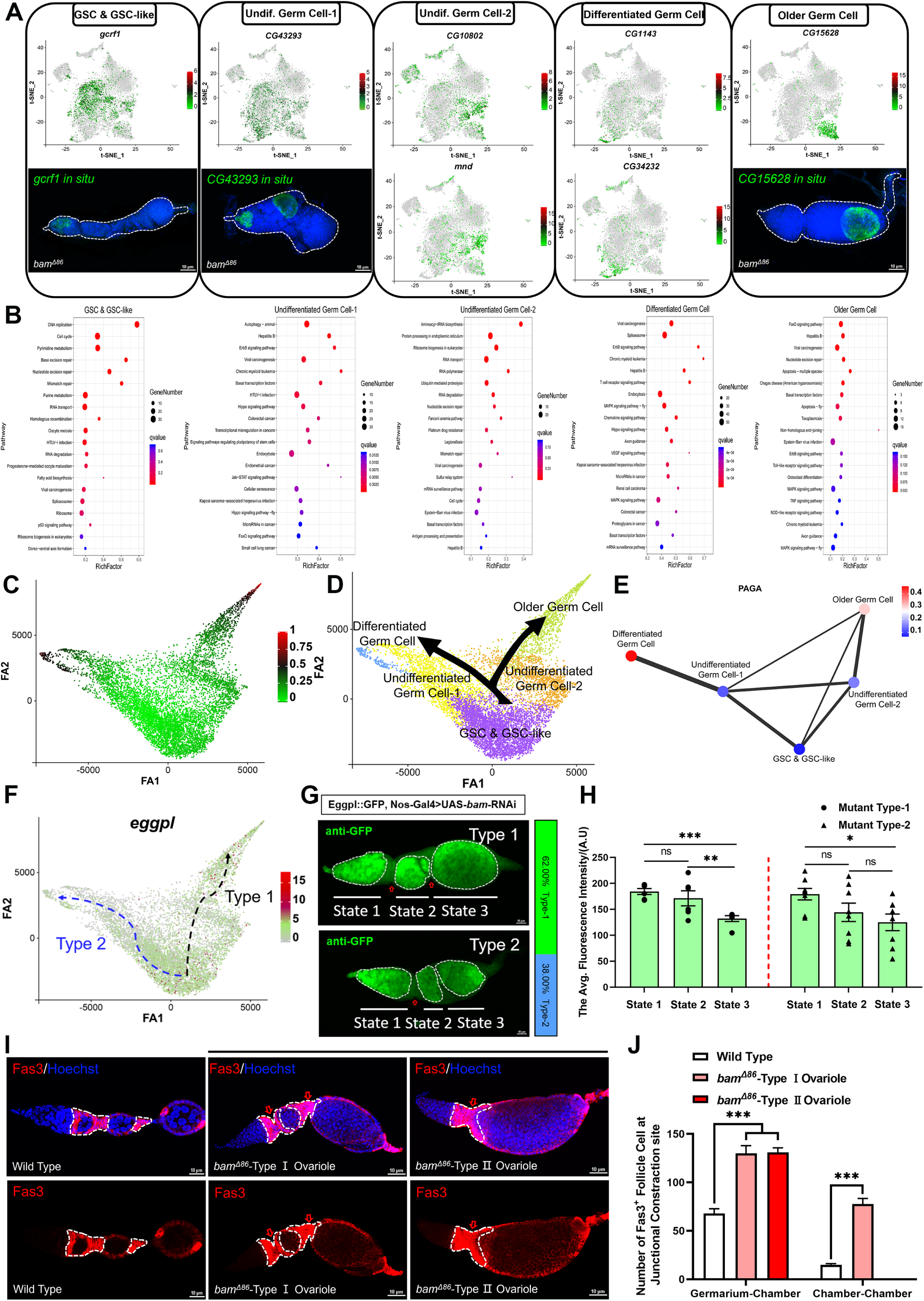
Identification of germline clusters and distinction of two specific *bam* mutant ovarioles. **(A)** Feature plots on tSNE shows the expression patterns of novel selected marker genes in distinct germline cell types in *bam* mutant ovary. *In situ* images of *gcrf1*, *CG43293* and *CG15628* **(B)** KEGG analysis on top 20 pathways of upregulated genes in 5 germ cell clusters. **(C)** PAGA-initialized single-cell embedding shows the predicted developmental trajectories of germ cell lineage. The bright green dots indicate the cells at early stage, while the red dots indicate the cells at late stage. **(D)** Annotation of cell types in PAGA-initialized single-cell embedding trajectories. Two arrowed lines indicate different developmental trajectories. **(E)** PAGA graph for visualized interaction among 5 germ cell clusters. The width of each lines represents the correlation intensity among these cell types. **(F)** Gene expression changes of *eggpl* along PAGA paths. **(G)** Two types of ovarioles in in *bam* mutant ovary labeled with anti-GFP. White dotted lines cycles 3 differentiation states, and red arrows points out the tissue constriction sites in two ovariole types. Type-1 ovariole account for 62% and the proportion of Type-2 ovariole is 38%. **(H)** The average fluorescence intensity of *eggpl*::GFP signals in Type-1 and Type-2 ovarioles. **(I)** Wildtype and *bam* mutant ovarioles stained with Fas3 (red) and Hoechst (blue). **(J)** Quantification of the number of somatic cells with Fas3^+^ signals at junctional constraction sites (germarium to chamber and chamber to chamber). Wilcoxon rank sum test is used as statistical analysis. All data are showed as mean ± standard deviation, and ns indicates no significant difference (*p* > 0.05), \**p* ≤ 0.05, \*\**p* ≤ 0.01 and \*\*\**p* ≤ 0.001.

To gain a deeper understanding of the distinct germ cell clusters in *bam* mutant germ cells, we used PAGA analysis (29) to infer developmental trajectories. As illustrated in Fig 3C-D, one transitional trajectory was from the GSC & GSC-like cluster to the Undifferentiated Germ Cell-1 and the Differentiated Germ Cell clusters, while the other was from the GSC & GSC-like cluster to the Undifferentiated Germ Cell-2 and Older Germ Cell clusters. Notably, the GSC & GSC-like cluster located at the beginning position was predicted to be a shared origin among two differential trajectories during tumor progression (Fig 3C-D). The PAGA graph also showed a close correlation among these two unique germ cell transitional trajectories (Fig 3E). To validate these inferred trajectories, we examined the expression pattern of *eggpl*, a marker for GSC and early cysts (23), in *bam^Δ86^*ovarioles (Fig. 3F). In *eggpl*::GFP, *nos*-Gal4>UAS-*bam*-RNAi lines, we observed 62% ovarioles showing a germarium and two chambers, and 38% of ovarioles with a germarium and one chamber (Fig. 3G). The fluorescence intensity of *eggpl*::GFP was quantified and indicated a significant difference of *eggpl* expression pattern between two types of ovarioles (Fig. 3H), suggesting a spatial heterogeneity or temporal variation of *eggpl* expression in *bam* mutant ovariole development. Further, Fas3 staining revealed abnormal accumulation of somatic cells at junctions of germarium to chamber in *bam^Δ86^* type Ⅰ and Ⅱ ovarioles and chamber to chamber in *bam^Δ86^* type Ⅰ ovarioles (Fig. 3I and 3J), highlighting disrupted follicle cells distribution and progressive structural disorganization in *bam* mutant ovaries.

### Comparison analysis uncovers transcriptomic characteristics of undifferentiated germ cells

To better understand the cellular heterogeneity of *bam* mutant ovary, we performed integration analysis of single-cell transcriptomic datasets from our wild-type and *bam*-mutant *Drosophila* ovaries, which revealed 19 distinct cell clusters with substantial overlap between datasets (Fig. 4A). Notably, the candidate marker *gcrf1* for early undifferentiated germ cell exhibited high expression in a specific population within cluster 0, and its expression was markedly increased in *bam*-mutant ovaries compared to wild type (Fig. 4B). Cluster-level expression analysis confirmed that *gcrf1*, together with the expression of germ cell markers such as *vas* and *eggpl*, was specifically enriched in cluster 0 (Fig. 4C). Subclustering analysis of cluster 0 further delineated five germline subpopulations (subclusters 0-4) (Fig. 4D). A comparison of cell composition revealed an overrepresentation of subclusters 0, 2 and 4 in *bam*-mutants, suggesting an accumulation of early undifferentiated germ cells (Fig. 4E). Differential gene expression analysis also predicted that both *gcrf1* and *eggpl* expression is highest in subcluster 0 and progressively decreases across the subclusters (Fig. 4F-G). SCENIC analysis revealed distinct differences in the regulon activity between wild type and *bam* mutant subclusters. Notably, Chrac-14, pho_extended and CG12391 exhibited elevated activity in subcluster ck_0, whereas their activity was markedly reduced in subcluster bam_0. Similarly, other regulons including Dref, mld, Stat92E_extended, nej and Dp showed a global shift from incresed activity in the wild type to reduced activity in the *bam* mutant group. By contrast, two regulons (Max and maf-S_extended) displayed higher activity in five subclusters from *bam* mutant. This result demonstrates that the silence of *bam* expression exerts a significant attenuation of transcriptional regulatory activity, thereby diminishing stemness and proliferative capacity in GSC and undifferentiated germ cells. (Fig. 4H). Furthermore, CellPhoneDB-based cell-cell communication analysis uncovered subcluster-specific interaction patterns, with both the number and strength of interactions varying across subclusters (Fig. 4I). Ligand-receptor pair analysis highlighted a cellular signaling pathway of NCAM1-related interactions in both wild-type and *bam*-mutant germline subpopulations (Fig. 4J), implying an important role in the developmental process of early undifferentiated germ cells.

**Fig. 4.**
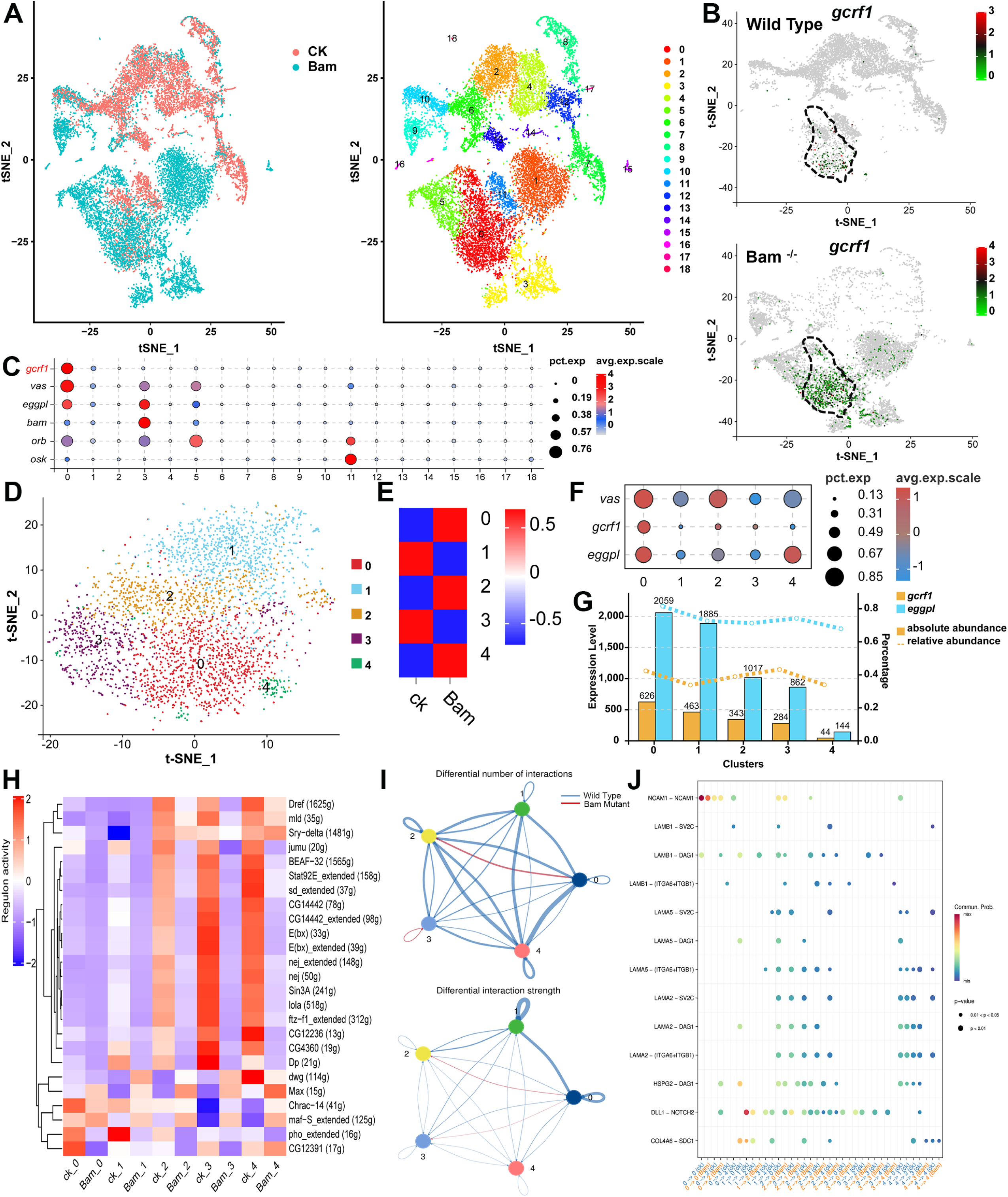
Integration of single-cell transcriptomic datasets identifies *gcrf1* as a marker for early germ cell and reveals characteristics of germline subclusters. (**A**) tSNE plot shows the overlap between cells from wild-type ovary dataset and cells from *bam*-mutant ovary dataset (left), and distinct 19 clusters in the integrated datasets (right). **(B)** The distribution of *gcrf1* expression in wild-type dataset and *bam*-mutant dataset. **(C)** Dot plot exhibits the enrichment of germ cell marker genes among 19 clusters. **(D)** tSNE plot highlights the 5 subclusters in the integrated cluster 0. **(E)** Comparison of cell quantity in 5 subclusters between wild-type dataset and *bam*-mutant dataset. **(F)** Dot plot shows the gene expression of *vas*, *gcrf1* and *eggpl* among 5 subclusters. **(G)** Changing expression level of *gcrf1* and *eggpl* from subcluster 0 to subcluster 4. **(H)** SCENIC analysis reveals different TF-based regulatory networks among 5 germline subclusters in both wild type and *bam* mutant. **(I)** Cycle plots of CellphoneDB analysis show the differential number of interactions (upward) and differential interaction strength (downward) among 5 subclusters. Different colors of outer circles represent distinct cell subpopulations, and the size of circle indicates the number of ligand-receptor pairs. The blue line indicates a stronger communication in the wild-type group, while the red line indicates a stronger communication in the *bam*-mutant group. The thicker line represents a greater communication strength. **(J)** Dot plot shows the ligand-receptor pairs contributing to different signaling among 5 subclusters in wild-type dataset and *bam*-mutant dataset.

### *gcrf1* is a specific regulator involved in germ cell development

We initially analyzed the conservation of Gcrf1 protein across different insect species by constructing phylogenetic tree. The result showed a strong similarity between the closest Gcrf1 homologs in *Drosophila* (78.26%-100%) and *Bactrocera* (58.59%-61.82%), and a weaker similarity to the homologous gene in *Culicidae* (50%) (Fig. 5A). This difference may reflect adaptive variations in gene function during the process of insect evolution. We validated the expression pattern of *gcrf1* by *in situ* hybridization and immunofluorescence staining. At mRNA level, *gcrf1* is specifically expressed in the anterior part of germarium where GSC and early cysts are located (Fig. 5B). In line with this, the HA signal in the *Gcrf1::HA* knock-in line overlapped with pMad and branched α-spectrin signals (Fig. 5C-D). Intriguingly, the expression pattern of *gcrf1* was restricted from the GSC to the 8-cell cyst in the *gcrf1* overexpression line triggered by *nos*-Gal4 (Fig. E). To investigate the role of *gcrf1* in GSC maintenance in the niche, we quantified the number of GSCs in *gcrf1* mutants. We found that RNAi knockdown of *gcrf1* specifically in germ cells with *nos-Gal4* caused a significant decrease in the number of GSCs per germarium whereas overexpression had the opposite effect, causing a significant increase in the number of GSCs per ovariole, which implies that *gcrf1* may be involved in GSC proliferation or the rate of GSC differentiation (Fig. 5F). Likewise, we found that ovary size was decreased upon RNAi knockdown of *gcrf1* and significantly increased upon overexpression of *gcrf1* in germ cells (Fig. 5G-H). In addition, we found that downregulation or upregulation of *gcrf1* could strikingly modulate the number of eggs laid per mated female (Fig. 5I). Lastly, we used MIST to examine the predicted interaction network of *gcrf1* and identified *CG15365* as a candidate. Indeed, we found by qRT-PCR that *CG15365* is significantly upregulated upon overexpression of *gcrf1* (Fig. 5J-K). Taken together, these results indicate that *gcrf1* is a novel regulator in the process of GSC renewal and germ cell development. Based on its expression in early germ cells and our finding that it is required for early germ cell function, we named this gene *germ cell renewal factor 1* (*gcrf1*).

**Fig. 5.**
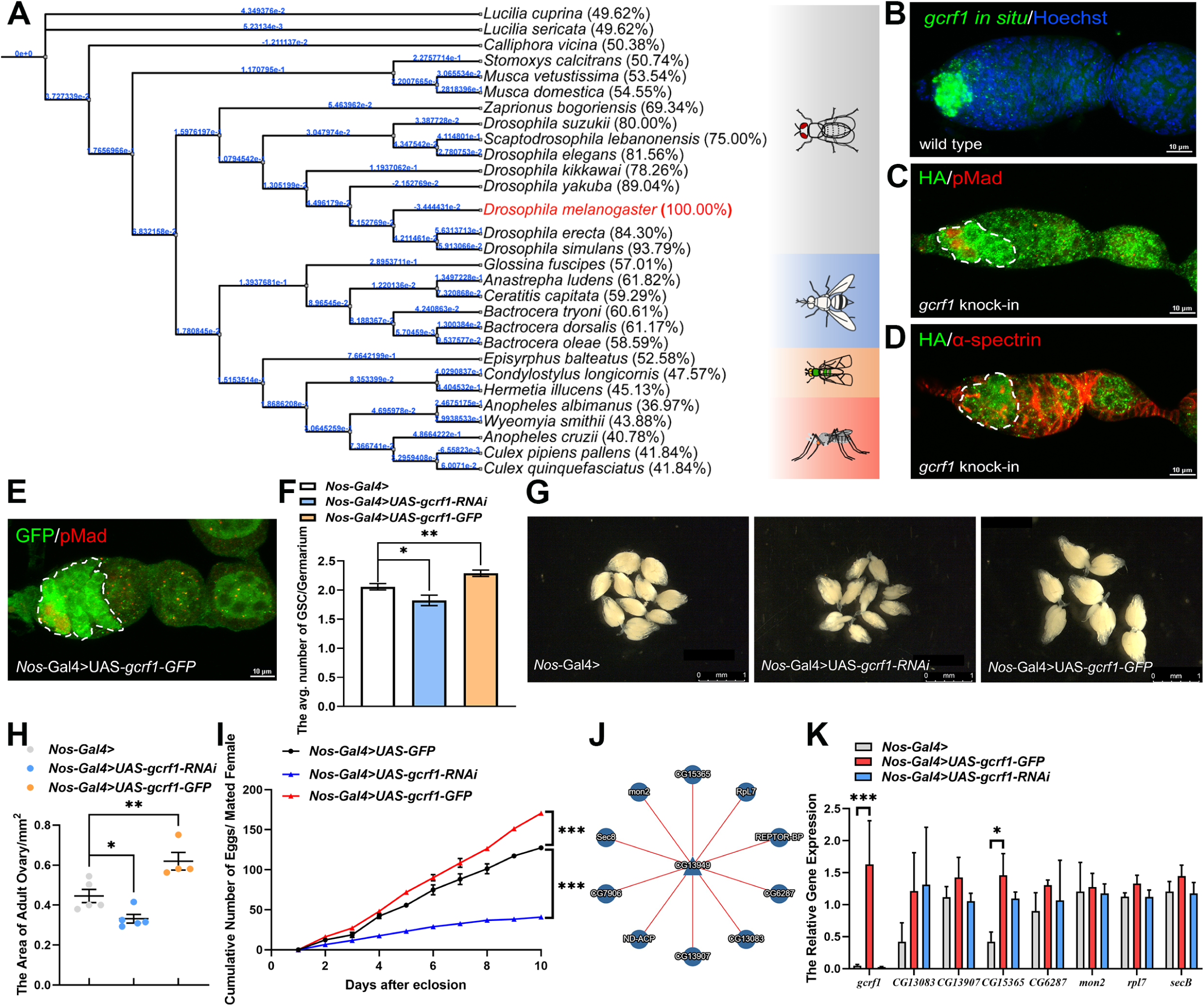
Functional analysis of *gcrf1* involving in female GSC proliferation and fecundity. **(A)** The construction of phylogenetic tree on gcrf1 protein conservation across species. **(B)** *In situ* hybridization of *gcrf1* (green) in wild-type ovariole labeling with Hoechst (blue). **(C)** The HA (green) and pMad (red) staining on *gcrf1* knock-in line. **(D)** The HA (green) and α-spectrin (red) staining on *gcrf1* knock-in line. **(E)** The GFP (green) and pMad (red) staining on *gcrf1* overexpression line triggered by *nos*-Gal4. **(F)** The average number of GSC in *nos*-Gal4, *gcrf1* RNAi and *gcrf1* overexpression lines. **(G)** The morphology of ovaries in *nos*-Gal4, *gcrf1* RNAi and *gcrf1* overexpression lines. **(H)** The quantification of ovary size in *nos*-Gal4, *gcrf1* RNAi and *gcrf1* overexpression lines. **(I)** The analysis of cumulative laid-egg numbers for each mated female flies over 10 days. **(J)** The prediction of PPI network based on Gcrf1. **(I)** RT-qPCR analysis on the potential interactive relationship according to PPI prediction.

## Discussion

In many cases, tumors are thought to originate from tissue-specific stem or progenitor cells that acquire genetic or epigenetic mutations, leading to uncontrolled growth. Additionally, cancer stem cells, a subpopulation within tumors, display stemlike properties and are often responsible for tumor initiation, metastasis, and resistance to therapy (31, 32). Bam is a master transcriptional regulator in the process of GSC differentiation, female flies with *bam* mutant ovary are sterile due to that all germ cells are arrested in an undifferentiated state, resulting in a failure of egg formation. A previous study demonstrated that two temporally distinct waves of gene expression occur during GSC differentiation, which includes an initial Bam-dependent shift primarily targeting cell cycle regulation and a reversion to a stem cell-like program that precedes terminal differentiation (33). In this work, we applied single-cell RNA sequencing to identify major cell types by using typical makers and decipher the cellular and transcriptional architecture of *bam* mutant *Drosophila* ovaries (Fig. 1B and C). Differential expression and PAGA-inferred trajectories show that germ cells split into two fates: one toward differentiation and one toward a senescence-like “older” state. This bifurcation demonstrates that loss of *bam* unlocks alternative developmental programs and mirrors observations that two undifferentiated subpopulations (Undifferentiated Germ Cell-1 and Undifferentiated Germ Cell-2) diverge from a common GSC and GSC-like origin (Fig. 3C-G). Undifferentiated Germ Cell-1 exhibits enrichment in mitotic and DNA-repair pathways, whereas Undifferentiated Germ Cell-2 shows enhanced protein processing and secretory signaling (Fig. 3B). These parallels with mammalian glioblastoma stem-like cells plasticity (34, 35) suggest that early tumorigenic lesions exploit both proliferative and stress-response programs to fuel growth. Notably, the activation of foxO and TNF pathways in older germ cells implies a programmed cell-death response that may serve as an intrinsic brake on unchecked proliferation.

Compared to prior scRNA-seq investigations of wild-type ovaries (23, 27, 36), our study is the first to characterize a GSC tumor model at single-cell resolution. The integration with published datasets via SC Transform confirms the robustness of our clusters and highlights the reproducibility of single-cell annotations across genotypes. The SCENIC regulon analysis further delineates transcriptional circuits underpinning these states: ovo activity predominates in the earliest undifferentiated subcluster, whereas Taf1 and crol regulons mark later transitional states (Fig. 4H). These findings resonate with previous single-cell studies identifying dynamic regulon shifts during germ cell differentiation, yet uniquely reveal how a single genetic lesion reshapes these regulatory landscapes to favor tumorigenic trajectories (37–39). Cell-cell communication analysis via CellPhoneDB uncovers subcluster-specific interaction networks, prominently featuring NCAM1-related ligand-receptor pairs in both wild-type and *bam* mutant (Fig. 4J). Given the role of NCAM in mediating cell adhesion and migration in cancer (40, 41), this suggests that aberrant germ cells coopt adhesion cues to sustain niche interactions and drive overproliferation.

Our discovery of *in vivo* validated marker *gcrf1* as a GSC-specific regulator that modulates proliferation and fertility represents a key mechanistic advance. Its specific expression pattern in GSC and early cysts, coupled with functional evidence that *gcrf1* knockdown reduces GSC number and ovary size, while overexpression of *gcrf1* has the opposite effects, highlights the essential role of *gcrf1* in female fecundity. Moreover, the high evolutionary conservation of *gcrf1* among *Culex*, *Bactrocera* and *Drosophila* species and its potential interaction with other germline-associated factors suggest it may serve as a key node in conserved stem cell regulatory networks.

We acknowledge several limitations of this study. First, given the operational complexity and limited sensitivity of *in situ* hybridization techniques, there are inherent technical constraints in their ability to resolve cellular heterogeneity within *bam* mutant *Drosophila* ovaries, thus incorporating spatial transcriptomic validation would strengthen the links between inferred developmental trajectories and their correspondence with cell populations. Second, although functional assays support the role of *gcrf1* in regulating GSC proliferation, its precise molecular mechanism underlying its property and function remain to be fully elucidated. Lastly, while *Drosophila* serves as a genetically tractable model, the extent to which *bam* mutant-driven tumorigenesis recapitulates features of mammalian germ cell tumors warrants further comparative investigation.

In conclusion, our single-cell atlas of *bam* mutant ovaries illuminate how GSC differentiation defects generate tumor-like overgrowth through the emergence of novel cell states, regulatory networks, and intercellular signaling circuits. This dataset thus serves as a foundational resource for dissecting reproductive tumorigenesis and future therapeutic explorations in reproductive and stem cell biology.

## Data availability

The raw sequence data reported in this paper have been deposited in the Genome Sequence Archive (Genomics, Proteomics & Bioinformatics 2021) in National Genomics Data Center (Nucleic Acids Res 2022), China National Center for Bioinformation / Beijing Institute of Genomics, Chinese Academy of Sciences (GSA: CRA028191) that are publicly accessible at https://ngdc.cncb.ac.cn/gsa. The referenced datasets in this study are available in GEO databases (GSE210822).

## Acknowledgments

This study was supported by National Key Research and Development Program of China (Grant No. 2023YFD1700700). We thank Ms. Jingjing Ning from Guangzhou Genedenovo Biotechnology Co., Ltd for the assistance in sequencing and bioinformatics analysis.

## Supplementary Figures and Tables

**Fig. S1.**
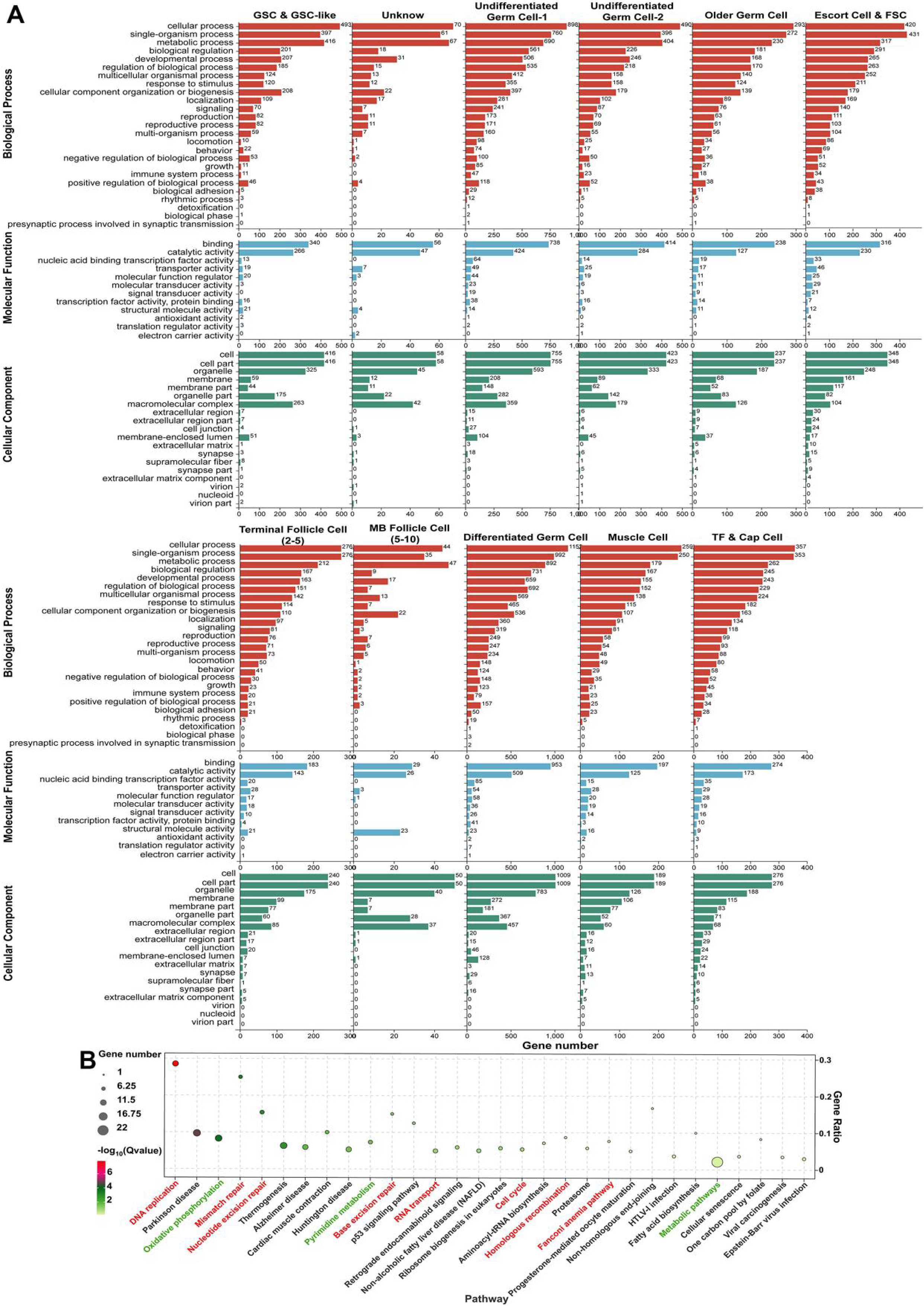
GO term analysis of all clusters. **(A)** A functional enrichment of 11 ovarian cells clusters annotated by GO database. **(B)** KEGG pathway analysis on the Unknown cluster. Pathways highlighted in red correspond to the major enriched pathways in mitotic follicle cell, whereas those in green denote the pathways enriched in endocycle follicle cells.

**Fig. S2.**
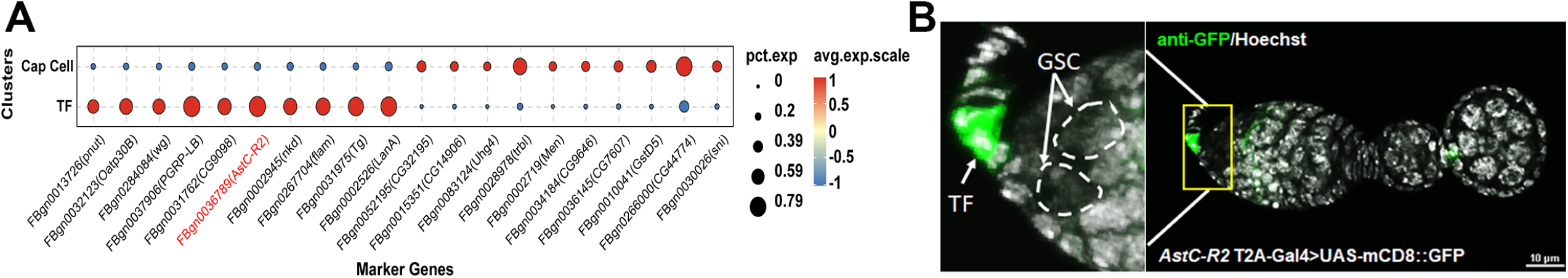
Identification of cap cell and TF cell cluster by *AstC-R2* expression. **(A)** Dot plot presents top 10 marker genes in Cap Cell and TF clusters respectively. **(B)** Anti-GFP (green) and Hoechst (white) staining on the ovary from *AstC-R2* T2A-Gal4>UAS-mCD8::GFP.

**Fig. S3.**
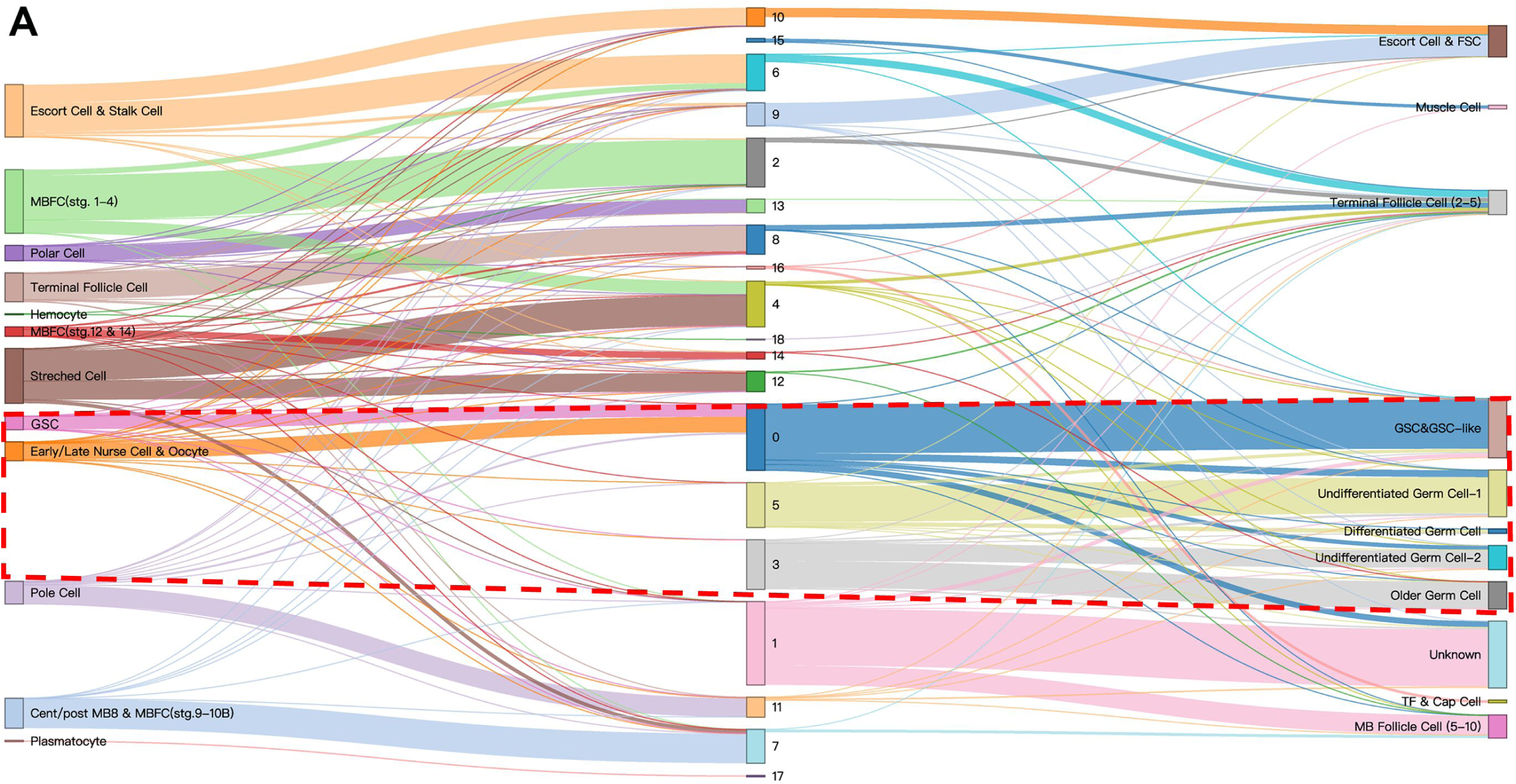
Identification of conserved cell types between wild-type ovary and *bam* mutant ovary. **(A)** Sankey graph shows the mapping between cell-type clusters in the *Drosophila* wild-type ovary (left) and cell-type clusters in the *bam^Δ86−/−^* ovary (right). The dotted line highlights the annotated germ cell types in wild-type and *bam^Δ86−/−^* ovary.

